# Deciphering the protein dynamics and molecular determinants of iPSC-derived neurons

**DOI:** 10.1101/599415

**Authors:** Suzy Varderidou-Minasian, Philipp Schätzle, Casper. C. Hoogenraad, R. Jeroen Pasterkamp, Maarten Altelaar

**Affiliations:** Biomolecular Mass Spectrometry and Proteomics, Bijvoet Center for Biomolecular Research and Utrecht Institute for Pharmaceutical Sciences, University of Utrecht, Padualaan 8, 3584 CH Utrecht, the Netherlands; Netherlands Proteomics Center, Padualaan 8, 3584 CH Utrecht, the Netherlands; Cell Biology, Department of Biology, Faculty of Science, Utrecht University, 3584 CH Utrecht, the Netherlands; Department of Translational Neuroscience, UMC Utrecht Brain Center, University Medical Center Utrecht, Utrecht University, 3584 CG Utrecht, the Netherlands

**Keywords:** iPSC, human, neuron differentiation, quantitative mass spectrometry, TMT-10plex

## Abstract

Neuronal development is a multistep process with different regulatory programs that shapes neurons to form dendrites, axons and synapses. To date, knowledge on neuronal development is largely based on murine data and largely restricted to the genomic and transcriptomic level. Advances in stem cell differentiation now enable the study of human neuronal development, and here we provide a mass spectrometry-based quantitative proteomic signature, at high temporal resolution, of human stem cell-derived neurons. To reveal proteomic changes during neuronal development we make use of two differentiation approaches, either by expression of neurogenin-2 (Ngn2) leading to glutamatergic induced neurons (iN) or via small molecule manipulations, leading to patterned motor neurons. Our analysis revealed key proteins that show significant expression changes (FDR <0.001) during neuronal differentiation. We overlay our proteomics data with available transcriptomic data during neuronal differentiation and show distinct, datatype-specific, signatures. Overall, we provide a rich resource of information on proteins associated with human neuronal development, and moreover, highlight several signaling pathways involved, such as Wnt and Notch.

## INTRODUCTION

The human brain is a complex system with different regions and cell types, having billions of cells and trillions of synapses [1, 2]. This diversity presents a great challenge in understanding the molecular and cellular function of this organ. As recent studies have revealed, perturbation of brain development underlies many neurological disorders such as autism and schizophrenia, however, much of our current knowledge is derived from the rodent brain [3–5]. Human neural development remains difficult to study given the ethical constraints in primary human brain tissues, together with the paucity of high-quality post-mortem tissue. Moreover, the degree of cell and tissue heterogeneity, in combination with complex developmental and environmental factors, further complicate (human) neuronal research [6, 7]. One approach with great promise to study neurological disorders is the use of induced pluripotent stem cells (iPSC) from e.g. human fibroblasts [8]. Ever since the first report on iPSC, major efforts have been directed towards developing differentiation protocols to induce neurons [9, 10]. Given the rapid developments in the field of iPSC-derived neurons, a comprehensive understanding of the mechanism underlying iPSC differentiation towards neurons is required. Neuronal development is coordinated by morphogens and neurogenic factors that can be captured *in vitro* using iPSC [11, 12]. Over the last years, major advances in iPSC differentiation improved the generation of homogeneous population of neurons and has been used to study various neurological disorders [13]. Although many of these regulatory pathways involved in neuronal development have been studied on genomic and transcriptomic studies, their mechanisms at protein levels have not [14]. Since proteins are the final molecular effectors of cellular processes and their perturbation is linked to pathological states, their investigation is essential.

Multiple protocols exist for generating neurons from iPSC. Here, to monitor the differentiation process of iPSC-derived neurons by high-resolution proteomics, we adapted two different approaches, often used to model neuronal development and neurological disorders [15–18]. Forced expression of a single neurogenic transcription factor (Ngn2) causes rapid differentiation of human iPSC into functional excitatory cortical neurons (iN cells) [19]. This approach shows, within 10 days, rapid and reproducible production of a homogeneous population of glutamatergic neurons. In addition, extrinsic-factor-based strategies of different morphogens such as Wnt, fibroblast growth factor (FGF), retinoic acid (RA) and sonic hedgehog (SHH) can be used to generate neuronal subtypes [20]. Here, the course of differentiation is a three-step process, with neural crest cell activation by dual SMAD inhibition, caudalization by RA signaling and ventralization by SHH signaling. We will refer to these neurons as patterned motor neurons (MNs). Both approaches can be used as model systems to study the molecular mechanisms during neuronal development.

The research presented here quantitatively probes proteome changes during differentiation of iN cells and patterned MNs at 10 different time points **(Figure 1)**. We observe a two-step resetting of the global proteome, showing abundant proteins in iPSC decreasing and neuronal proteins increasing over time. We highlight both well-established and novel proteins up- and downregulated during differentiation. Additionally, we compare our proteomic data with comparable transcriptomic data during neuronal differentiation showing distinct (gene/protein) signatures, and we illustrate the relative fold change of proteins associated with signaling pathways such as Wnt, Notch and hedgehog signaling. Finally, we illustrate which proteins are specifically changing during differentiation of iPSC into either iN cells or patterned MNs.

**Figure 1.**
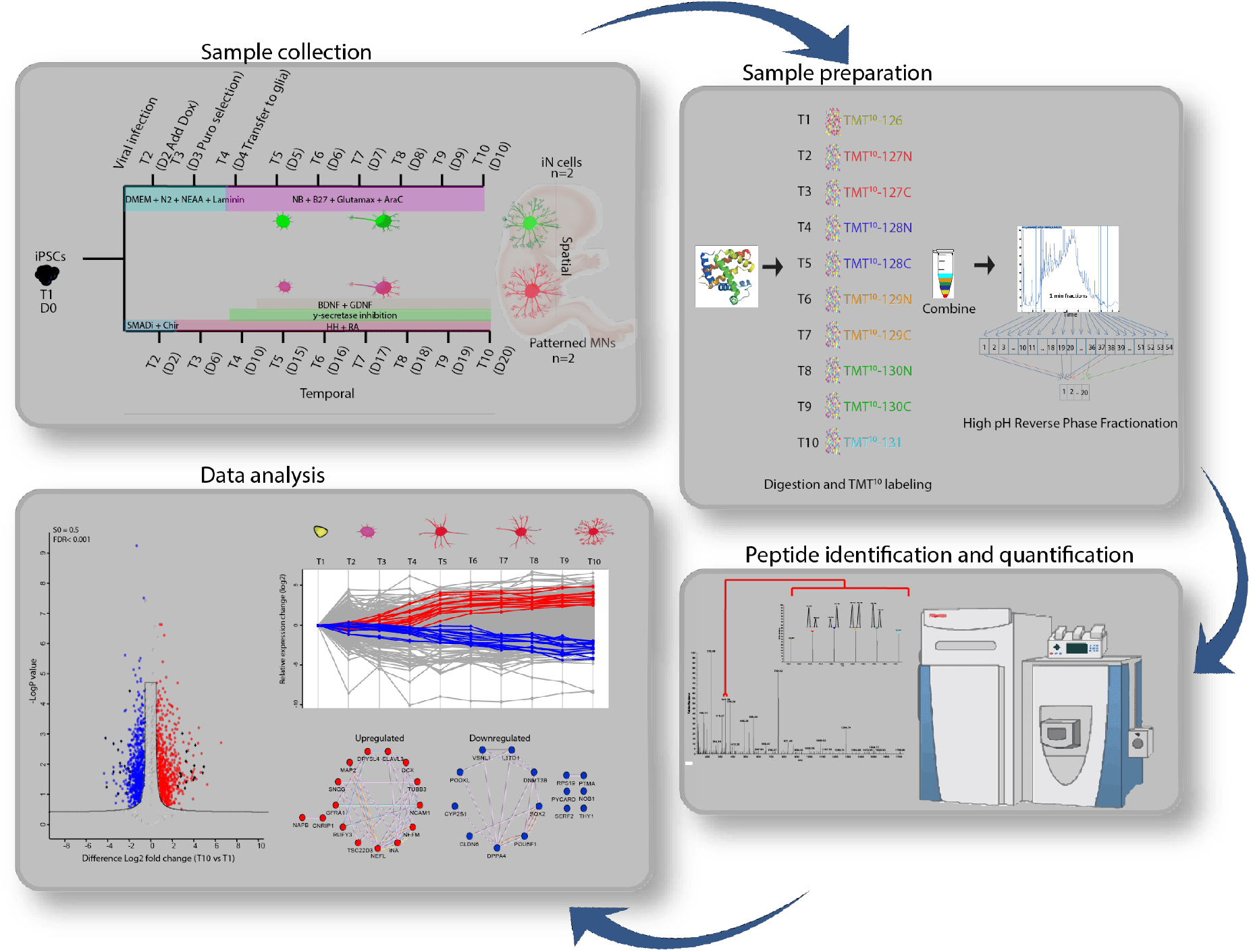
Workflow of MS-based quantitative proteomics during neuronal differentiation. Differentiation of iPSC towards iN cells was performed using doxycycline-induced expression of Ngn2. Differentiation of patterned MNs was performed using the action of small molecules for neural induction and cell fate determination. Proteins extracted at ten time points from two biological replicates for each cell type were digested and TMT 10-plex labeled. Peptides were combined and fractionated using high pH fractionation. The resulting fractions were analyzed by high-resolution nano-LC-MS/MS and quantification was achieved using TMT 10-plex isobaric labeling.

## RESULTS

### Differentiation towards iN cells and patterned MNs

Differentiation into iN cells was performed by doxycycline-induced expression of neurogenin-2 (Ngn2) [19]. At day 7, the iPSC produced an apparently mature neuronal morphology and were positive for the neuronal marker TuJ1 and the cortical marker FOXG1 (**Figure 2A**). In addition, these cells were negative for MN marker ISLET1. Patterned MNs were generated by the combined actions of small molecules for neural induction and cell fate determination [20] **(Figure 2B, supplementary video)**. At day 20, cells were positive for βIII-tubulin (>80%) and ISL1 (~40 %) markers. For mass spectrometry-based quantitative proteomics, we performed TMT-10plex labeling of the tryptic peptides originating from 10 distinct time points within the differentiation timeline of either iN cells or patterned MNs, resulting in a total of four biological replicates **(Figure 1)**. After separation by high-pH fractionation, the labeled peptides were analyzed with high-resolution tandem mass spectrometry (LC-MS/MS).

**Figure 2.**
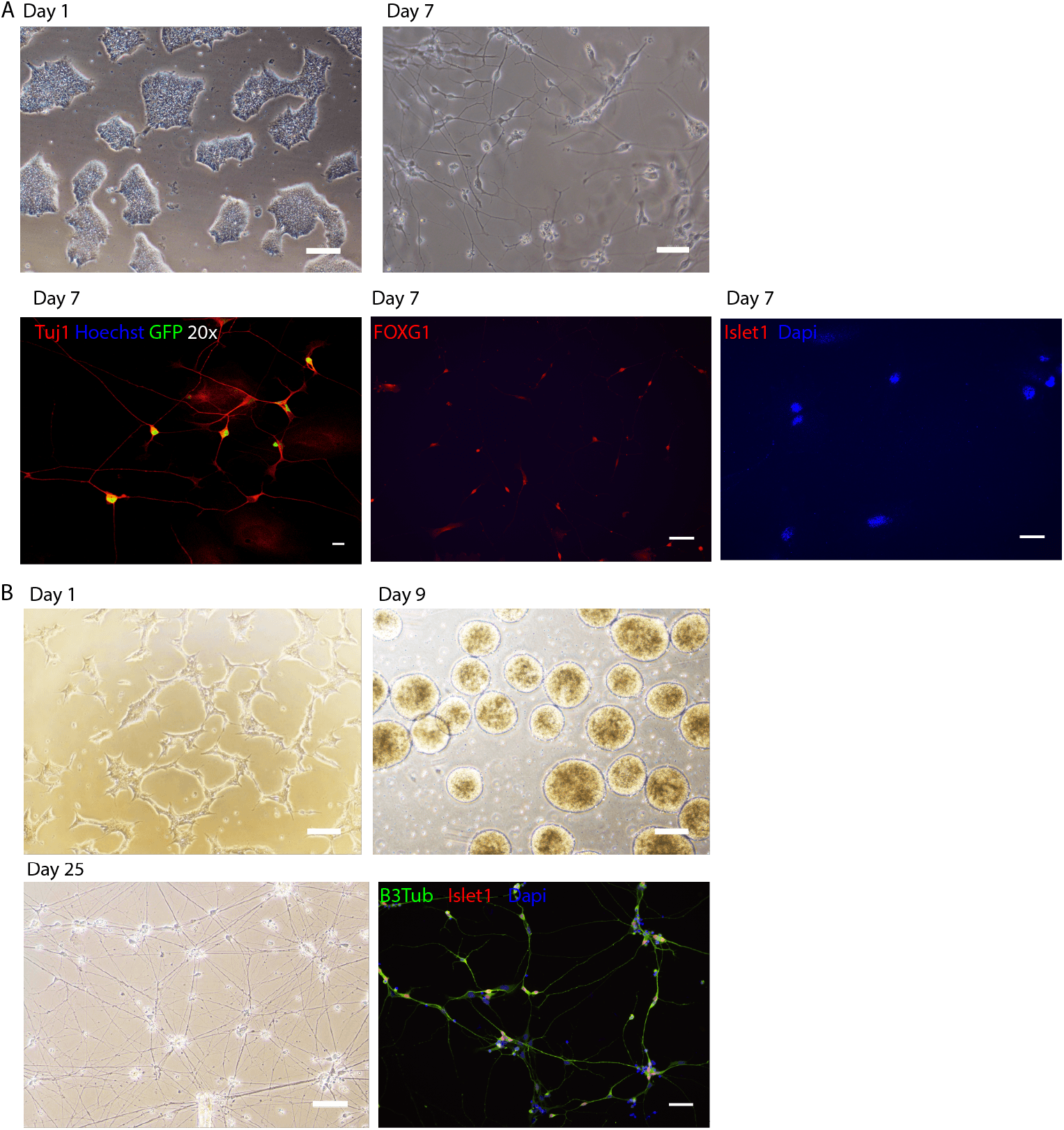
Bright field images and immunocytochemistry of iN cells and patterned MNs at indicated time points. (A). Bright field images of iPSC at day 1, iN cells at day 7 and immunostainings for neuronal marker (TuJ1), cortical marker (FOXG1) and motor neuron marker (Islet1). (B). Bright field images of patterned MNs and immunocytochemistry staining for neuronal marker (Tubulin beta 3), motor neuron marker (Islet1) and nuclear marker (Dapi). Scale bars = 100 μm

### MS-based quantitative proteomics

We identified 4070 and 3517 protein groups with a false discovery of 1%, in iN cells and in patterned MNs, respectively **(Figure S1A)**. The relative abundances of the quantified proteins span more than four orders of magnitude, indicating a broad dynamic range in our quantitative measurement. Tissue enrichment analysis of the 25% most abundant proteins against the whole human proteome background, revealed enrichment for ‘Cajal-Retzius cell’, ‘fetal brain cortex’ and ‘epithelium’ **(Figure S1B)**. The expression of Cajal-Retzius cells is not surprising since these cells are involved in the organization of brain development [21]. The enrichment associated with epithelium may be caused by origin of the fibroblast. Finally, annotation of protein class for all identified proteins revealed a wide coverage, including typically low abundant protein classes such as transcription factors and storage proteins **(Figure S1C)**. Furthermore, high run-to-run quantitative reproducibility was observed between biological replicates both for iN cells and patterned MN differentiation **(Figure S1E)**. The TMT reporter intensity values of proteins identified in each time point were normalized to the reference intensity value of these proteins before initiation of differentiation (T1). As a consequence, all data throughout the study reflects a ratio change relative to T1. After global analysis of our proteomics data, we set out to confirm the quality of the neuronal differentiation. We investigated known stem cell markers (e.g. SOX2 and OCT4) that indeed decreased during differentiation while, as expected, several neuronal markers (e.g. NEFL, GAP43 and MAP2) increased in both neuron types **(Figure 3A)** [22–28]. Progenitor markers (e.g. BIRC5, SPARC and TOP2A) that are required for fetal development and regulation of cortex development show, in our data, higher expression in iN cells compared to patterned MNs [29–32]. Furthermore, the rostral marker OTX1 was only identified in iN cells and the caudal marker HOXB5 only in patterned MNs. Finally, NEUROG2 expression was only observed in iN cells, where it is overexpressed, and previous studies showed its involvement in cortical specification [33]. Taken together, these data confirm the validity of our approach, showing neuronal subtype-specific development.

**Figure 3.**
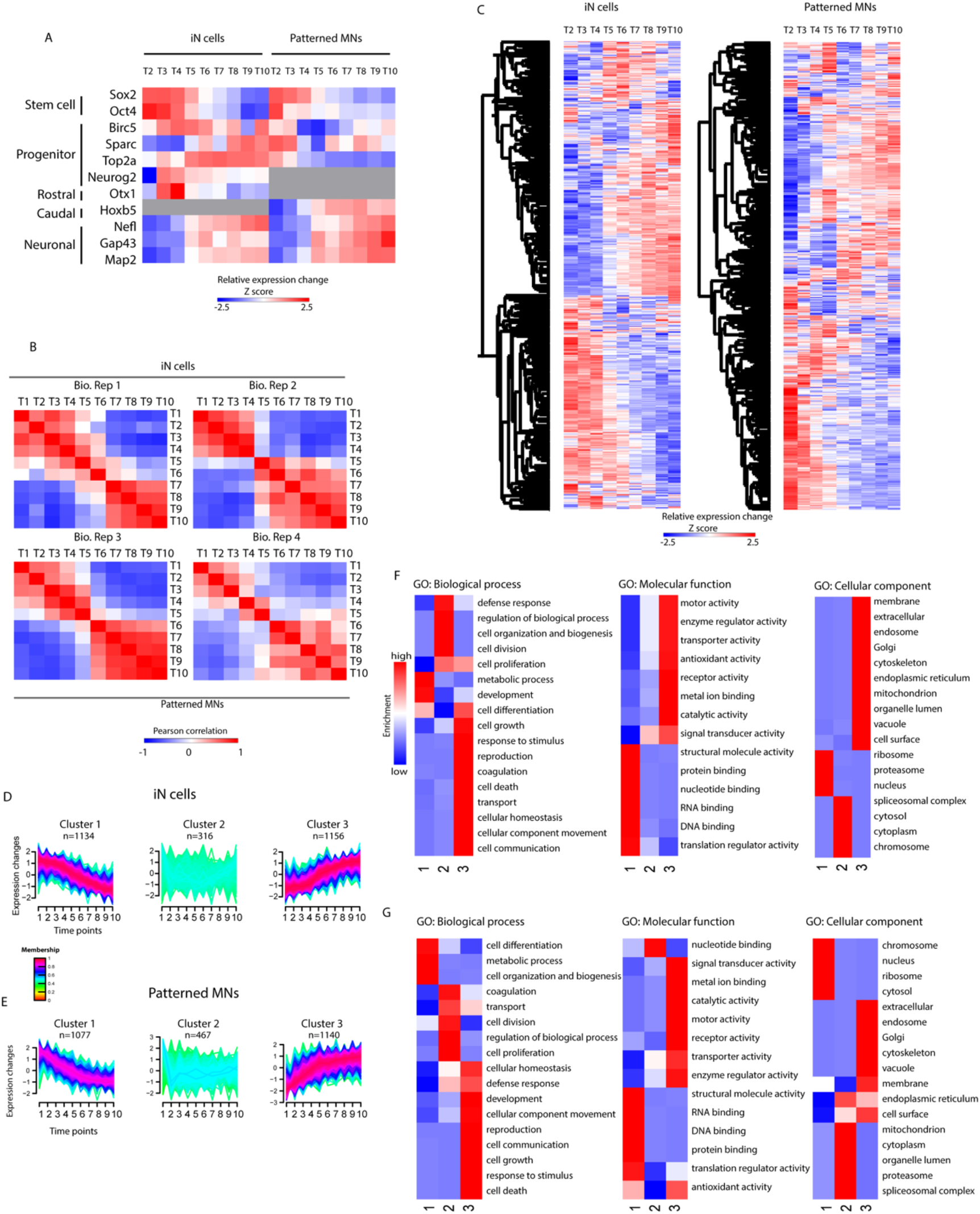
Proteome dynamics during neuronal differentiation. (A). Differential expression of general iPSC, progenitor and neuronal markers represented in a heat map. Each row represents average protein log2 ratios relative to iPSC (T1) in both replicates for each approach. (B). Pearson correlation within each biological replicate. Note for each biological replicate the high correlation between early time point (T1-5) and late (T6-T10). (C). Heat map of all proteins identified in iN cells and patterned MNs showing the expression changes along the course of differentiation for each approach relative to T1. (D-E). Clusters of the protein dynamics during differentiation towards iN cells (E) and patterned MNs (E). Three clusters with increasing, decreasing and not changing profiles were revealed in both differentiation approaches. Upper and lower ratio limit of log2 (0.5) and log2 (−0.5) was selected for inclusion into a cluster. The number of proteins is represented by n while the membership value indicates how well each protein fits to the average cluster profile. (F-G). Gene ontology enrichment analysis of each cluster tested for overrepresented biological processes (BP), molecular function (MF) and cellular component (CC) compared to unregulated proteins.

To obtain a global proteome view, changes of proteins along the course of differentiation for the two approaches were evaluated **(Figure 3B, C)**. We observed a ‘two-step resetting’ of the global proteome, with proteins highly expressed in iPSC showing down-regulation over time, and vice versa, proteins with low expression in iPSC increasing over time. This distinct separation of the proteome, which happens around the intermediate stage (T5), independent of the differentiation approach, can be further emphasized by performing Pearson correlation within each biological replicate (**Figure 3B**). Here, an anticorrelation (R = between −0.58 and −0.83) was found between T1 and T10, which was similarly observed during reprogramming of somatic cells to pluripotency [34]. Next, we performed an unsupervised clustering of the protein expression across the time points for both differentiation approaches **(Figure 3D, E)**. Gene Ontology (GO) enrichment analysis of these clusters **(Figure 3F, G)** revealed biological processes such as cell communication, growth and movement being upregulated (cluster 3), while cell differentiation and biogenesis is slowly downregulated (cluster 1). Closer inspection of the corresponding proteins in cluster 3 revealed pan-neuronal proteins such as tubulin-beta-3, neurofilament and MAP2. Cluster 3 also reveals increased expression of proteins localized in the membrane, cytoskeleton and several organelles, as well as molecular functions related to transport, enzymatic activity and signaling. Cluster 1 contains nuclear, chromosomal and ribosomal proteins that gradually decrease over time having a function in RNA and DNA binding. Downregulation of RNA processing during differentiation was previously shown in mouse by proteomics analysis and in human by transcriptomic analysis [14, 35]. Cluster 2 includes a group of proteins with general terms such as regulation of biological processes or cell division.

### Proteomic changes across time points during neuronal differentiation

To identify differentially expressed proteins during neuronal differentiation in the two approaches, we compared T10 relative to T1 and performed a t-test for both neuronal subtypes separately. **Figure 4A, C** show the volcano plots with the most extreme comparison T10 (mature neurons) against T1 (iPSC) for both approaches, depicting proteins with a FDR below 0.1%. This resulted in 696 significantly downregulated proteins and 686 upregulated proteins in iN cells and 706 significantly downregulated proteins and 620 upregulated proteins in patterned MNs **(Supplementary Table 1, 2)**. Close inspection of the proteins significantly enriched in T10 reveals general microtubule-associated proteins necessary for neuronal stabilization, outgrowth and migration, whereas specific iPSC enriched proteins consisted of transcription factors and proteins having a role in embryonic development. Furthermore, we highlight the top 15 proteins showing the largest fold change over time in both neuronal differentiation approaches **(Figure 4B, D)**. As expected, proteins such as SOX2, POU5F1 (OCT4) (iPSC markers) and MAP2 and DCX (neuronal markers) were enriched. Moreover, several proteins (e.g. INA, DPYSL4, and RPS19) were identified, that have less established roles in the context of pluripotency and differentiation [36–38]. The analysis of these regulated proteins shows an interaction network around the proteins INA, NCAM, NEFM and NEFL, being upregulated, and POU5F1, SOX2, DNMT3B and DPPA4, being downregulated **(Figure 4E)**.

**Figure 4.**
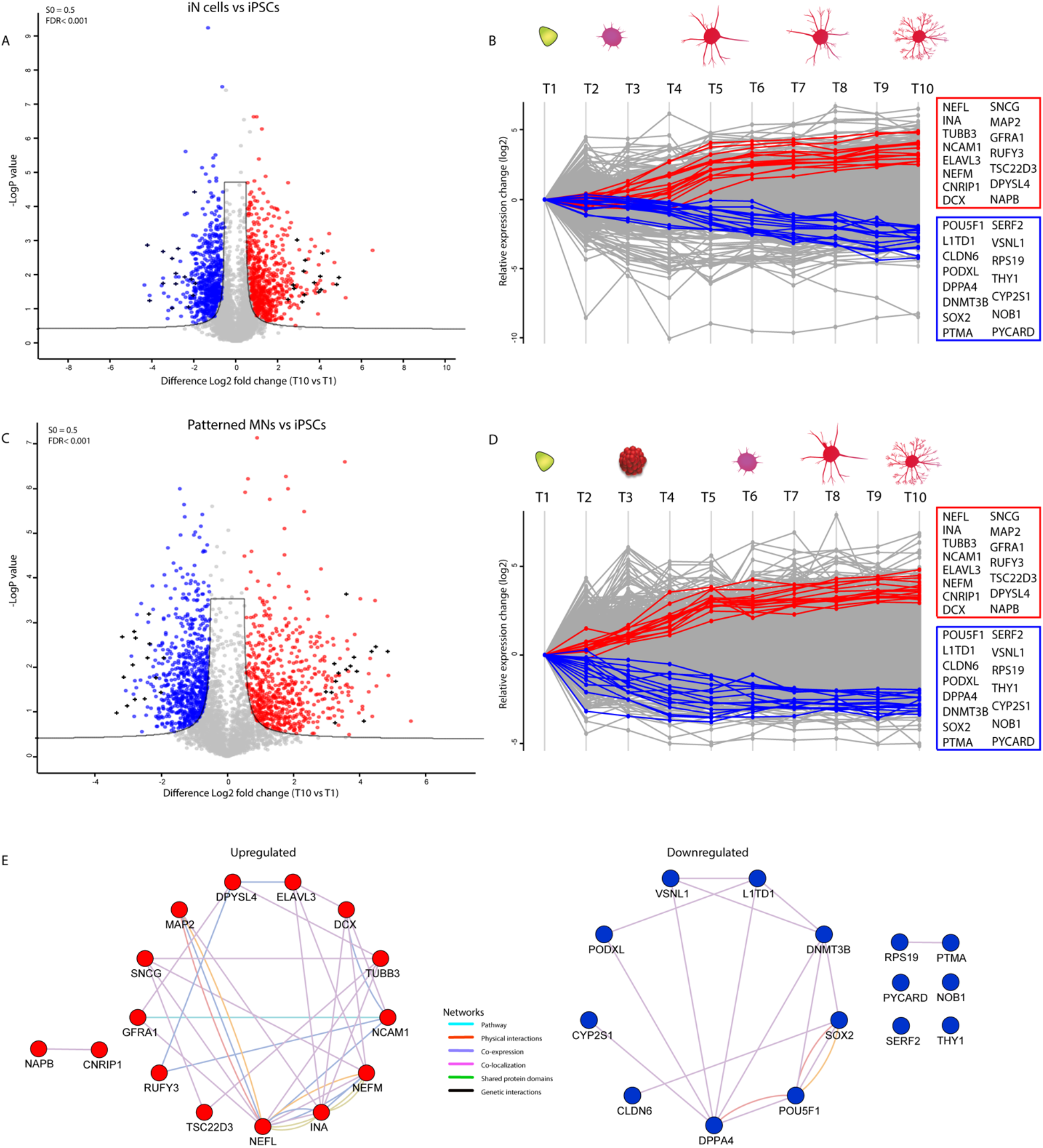
Significance analysis of protein expression changes during neuronal differentiation. (A) Volcano plot illustrating differentially expressed proteins during iN cell differentiation. The –log10 P value is plotted against the log2 fold change of T1 (iPSC) and T10 (mature iN cells) with significant threshold t-test with FDR < 0.001 and S = 0.5. Red represents proteins up regulated in T10 and blue represents proteins downregulated in T10. Black cross represents the top 15 proteins with the highest fold change. (B) Profile plot highlighting the top 15 significant up- and downregulated proteins in iN cells. (C) Volcano plot illustrating differentially expressed proteins during patterned MN differentiation. The –log10 P value is plotted against the log2 fold change of T1 and T10 with significant threshold t-test with FDR < 0.001 and S = 0.5. Red represents proteins up regulated in T10 and blue represents proteins downregulated in T10. Black cross represents the top 15 proteins with the highest fold change. (D) Profile plot highlighting the top 15 significant up- and downregulated proteins in patterned MN differentiation. (E) Protein network analysis showing the top 15 upregulated or downregulated proteins that interact with each other.

Next, we compared the significantly altered proteins during differentiation towards iN cells or patterned MNs, and observed only moderate overlap (23%) (**Figure 5A**). The differences in protein expression might suggest specificity towards a neuronal subtype or differentiation approach. We selected proteins with the largest fold change between the two neuronal subtypes (fold change cut-off ≥ 1.5 and p value ≤ 0.1). Proteins that increase most in iN cells (compared to patterned MNs), are S100A11, S100A13 and S100A6 **(Figure 5B)**. The S100 family of calcium binding proteins was first identified in the brain, regulating processes such as cell cycle progression and differentiation [39]. They have been found in sub-cortical structures and in a subpopulation of astrocytes [40–43]. Proteins highly enriched in patterned MNs are PPARD and MDN1. They act downstream of the Wnt-β-catenin and NOTCH signaling pathway [44, 45]. We then selected proteins exclusively identified either in iN cells or patterned MNs that show a minimum twofold change in at least one time point during differentiation **(Figure S5)**. The highly expressed protein SULF2 in iN cells is involved in brain development and neurite outgrowth in mice [46, 47] but its involvement in human neuronal development is unclear. Furthermore, highly expressed proteins in patterned MNs (e.g. PBX3 and HOXB5) regulate dorsal spinal cord development which is in line with the neuronal origin [48].

**Figure 5.**
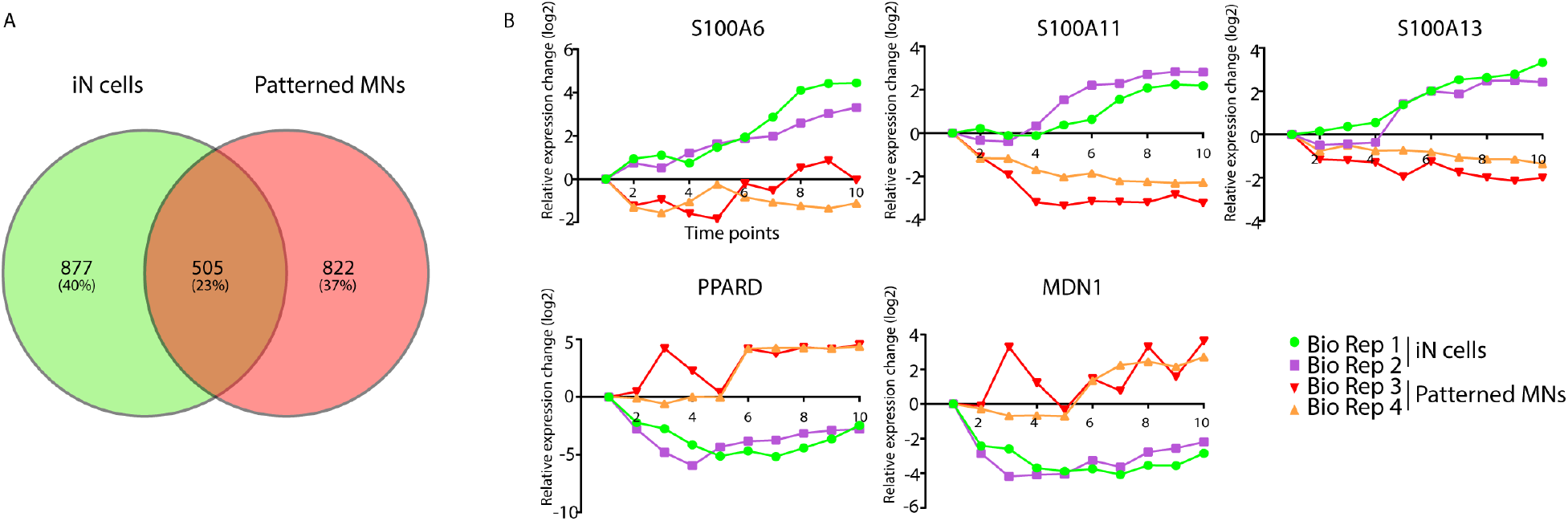
Differential expressed proteins in iN cells and patterned MNs. (A). Venn diagram indicating the overlap between the differentially expressed proteins in iN cells or patterned MNs (FDR 0.1%). (B) Example of proteins showing the largest differences between iN cells and patterned MNs during differentiation.

### Neurogenesis associated proteins

To further investigate proteins involved in specific processes related to neuronal differentiation we next looked for enrichment of proteins associated with neurogenesis using the DAVID database (https://david.ncifcrf.gov/summary.isp). This analysis revealed 129 proteins for which we had a closer look in their differentiation profile, plotting their expression over time in a heatmap **(Figure S2)**. These neurogenesis-associated proteins for the largest part increase their expression over time, such as DCX, CPLX2 and ELAVL3. Upregulated microtubule associated protein DCX has been increasingly used as a marker for neurogenesis and ELAVL3 is a neuron-specific protein [49, 50]. Several proteins also show downregulation, such as SLC7A5 and the transcription factor ZIC2, known to play a role in neuronal cell proliferation (neurogenesis) and early stages of central nervous system development [51, 52]. Additionally, ARF6 is known to regulate neuronal development in the brain via regulation of actin dynamics and synaptic plasticity. Its relative lower expression here however, might indicate that additional factors are needed to fully recapitulate the neuronal development.

### Transcription factors and cytoskeletal proteins

During neuronal differentiation, we showed that most of the up- and downregulated proteins are cytoskeletal and transcription factors (TFs), respectively. To better categorize the involvement of these proteins in neuronal development, we aimed to capture all TF’s and cytoskeletal proteins identified in our proteomic data **(Figure S3)**. TFs play essential roles in both reprogramming and neuronal development and have been used to induce neuronal differentiation from multiple cell lineages such as human fibroblasts, neuronal progenitors and stem cells [53–55]. We identified 259 regulated TFs (with a fold change cut-off ≥ 2 in at least in one time point for both approaches separately) of which PHC2, CRTC1, NCOA3 and SMAD3 are the most upregulated in both neuronal differentiation approaches **(Figure S3A)**. Interestingly, PHC2 has been reported to have an early developmental role in Drosophila, while its specific function in human cells has not been determined. CRTC1 is ubiquitously expressed in fetal brain and acts as a coactivator of the CREM (cAMP-responsive element binding)-dependent gene transcription pathway [56]. Furthermore, CRTC1 has been associated to different neuronal functions, such as synaptic plasticity and dendritic growth in cortical neurons and is also down-regulated in Alzheimer’s disease [57, 58]. NCOA3 has been identified as a novel microRNA regulator of dendritogenesis in mouse cortical neurons [59]. In line with this, we show that NCOA3 increases over the course of human neuronal differentiation. SMAD3 which was shown to promote neuronal differentiation in spinal cord of zebrafish [60] was found to be higher expressed in patterned MNs compared to iN cells. In addition, several TFs are upregulated at early time points of neuronal differentiation. NFXL1 was upregulated at the early time points of both neuronal subtypes while CCNT1 and LITAF for iN cells and ZIC5, NFATC4 and ZBTB40 for patterned MNs. Both ZIC5 and NFATC4 are essential in neural development and survival, however their association in early stages of caudalization, as seen here, has not been studied yet [61, 62].

In addition, we captured 256 cytoskeletal proteins as shown in **Figure S3B** with TUBB3, TAGLN3 and MAP1LC3A being most upregulated in both differentiation approaches. TUBB3 and MAP1LC3A are highly expressed in neurons and function in neurite formation and stabilization [63, 64]. TAGLN3, is an actin-binding protein involved in cytoskeletal organization [65]. Interestingly, the transgelin family members TAGLN, TAGLN2 and TAGLN3 show differential expression between iN cells and patterned MN differentiation. **(Figure S3C)**. TAGLN decreases during patterned MN differentiation while in iN cells there is an increase only in the early time points. TAGLN2 has a moderate increase in iN cells and decreases in the early time points of patterned MN differentiation. Only the third member of the transgelin family (TAGLN3) shows strong upregulation during neuronal differentiation in both approaches.

### Signaling pathways

In the last 30 years, several studies have demonstrated the importance of signaling pathways for regulating the developmental program, specify cell fate and patterning [66, 67]. Some of these signaling pathways will be discussed here, displaying the expression of their associated proteins in heatmaps for both iN cells and patterned MNs **(Figure S4)**. The TGFβ/ BMP signaling is known to inhibit neuronal development by blocking the proliferation of precursor cells in the adult brain. The dual SMAD inhibitor LDN-193189 and SB-431542 (together SMADi) induce the specification of cells with neural plate identity by selectively inhibiting the TGFβ/ BMP signaling pathway [68]. We applied this to our patterned MN differentiation protocol (at T2) and identified expression changes for all SMAD members, as well as SMAD interacting protein 1 (SNIP1) over time **(Figure S4A)**. SMAD3 rapidly increases at the late time points of patterned MN differentiation whereas the other members remain more constant. Interestingly, we observed in our iN cell differentiation protocol, that did not use of SMADi a similar expression pattern over time for most of the SMAD family members. Wnt signaling regulates cell migration, cell polarity and neuronal development during embryonic development and has an essential role in body axis formation, particularly in the formation of anteroposterior and dorsoventral axes [69–71]. Furthermore, the combinatorial effects of Wnt, retinoic acid (RA) and hedgehog (HH) signaling specify neuronal subtypes [72, 73]. Upon activation of Wnt signaling in patterned MNs (by Chir-99021 at T2 for four days), elevated levels of WNT5A, WLS, ZBTB16 and NFATC4 are observed in the early time points of patterned MNs while not identified or very low expressed in iN cells **(Figure S4B)**. These proteins are involved in cell fate pattering and development [74–77]. PLCB1 and PLCB4 are highly expressed in iN cells, compared to patterned MNs. A previous study showed their high expression and differential distribution in several regions of the brain including cortex, suggesting a specific role in different regions of the brain [78]. Proteins associated with hedgehog (HH) and retinoic acid (RA) signaling act as posteriorizing agent for spinal cord development [79, 80]. Both HH and RA are induced at T3 in the patterned MN differentiation protocol and kept till the end of the experiment. We looked for proteins associated with HH and RA and observed that the majority of these proteins are slightly elevated in patterned MNs compared to iN cells **(Figure S4C, D)**. CSNK1E and GSK3B together with CCND2 protein are higher in expression in the patterned MNs compared to iN cells. GSK3B has a critical role in axonal regulation and CCND2 has a function in neuronal development [81, 82].

The expression of cellular retinoic acid-binding protein 1 and 2 (CRABP1 and CRABP2) is increased in the patterned MNs after addition of RA. Interestingly, CRABP1 expression was restricted to certain neuronal subtypes in the hypothalamus suggesting for a cell-type specific function in the brain [83]. Homeobox proteins, which promote the expression of posterior neural genes, are all identified in the patterned MNs but not in the iN cells **(Figure S4E)** [84].

Inhibition of Notch signaling is induced (at T4) for the patterned MN differentiation, which is necessary to induce proneural genes [85, 86]. We identified 81 proteins involved in notch signaling, of which THBS1 showed a direct response after induction in the patterned MNs **(Figure S4F)**. THBS1 has been shown to play roles in neuronal development, such as in neurite outgrowth and cell migration [87, 88]. We further explored our data for enrichment of axon guidance proteins. They are important in regeneration along the anterior-posterior axis [89]. We identified 36 proteins associated with axon guidance of which the majority increases along the course of patterned MN differentiation **(Figure S4G)**. As described previously, axonal guidance cues are often categorized as ‘attractive’ or ‘repulsive’ [90]. NTN1 and its receptor (DCC), having attractive roles, are both increased in expression only in patterned MNs whereas ENAH and VASP, which function downstream of the repulsive axon guidance receptor Robo, are downregulated along the course of differentiation for both approaches [91].

### Comparison of neuronal development expression profile by quantitative proteomics and transcriptomics

We compared our proteomic measurements with RNA-seq data derived from a comparable transcriptome analysis during neuronal differentiation by overexpression of two transcription factors (Neurogenin-1 and Neurogenin-2) that resulted in homogeneous population of neurons within 4 days [14]. The RNA-seq data covers four time points with T1, which resembles the iPSC, and T4, which resembles mature neurons across the full course of iPSC differentiation. Out of 16589 transcripts that were identified, 5010 proteins were identified in our proteomics data. We selected the top 15 up- or downregulated transcripts and proteins during the differentiation and compared their expression levels (**Figure 6**). Interestingly, the majority of the transcripts that were downregulated showed a relative high protein expression level, as for instance TPD52L1, AKR1C3 and DAB2. Recent data have shown that these proteins are regulators of cell differentiation [92–94]. The majority of these most upregulated transcripts were upregulated in proteomics data for both neuronal subtypes and, vice versa, the majority of these upregulated proteins were upregulated in transcriptomic data. Interestingly, DCX, ELAVL3, GFRA1 and PRPH were identified as among the top 15 most upregulated (transcripts/proteins) in both transcriptomic and proteomic analysis separately, further emphasizing their crucial role in neuronal differentiation. They are all expressed in mature neurons and adult brain, however, only DCX is used as a neuronal marker [95–98]. The majority of the most downregulated proteins decrease to a lesser extent in transcriptomic data. PTMA for example, which is only downregulated on proteome level, was previously shown to be highly expressed in iPSC and is one of the validated targets of the c-MYC reprogramming transcription activator [99]. Together, these variations, between transcriptome and proteome, suggest that additional mechanisms such as posttranscriptional regulatory mechanisms might control the expression level of several proteins involved in neuronal differentiation.

**Figure 6.**
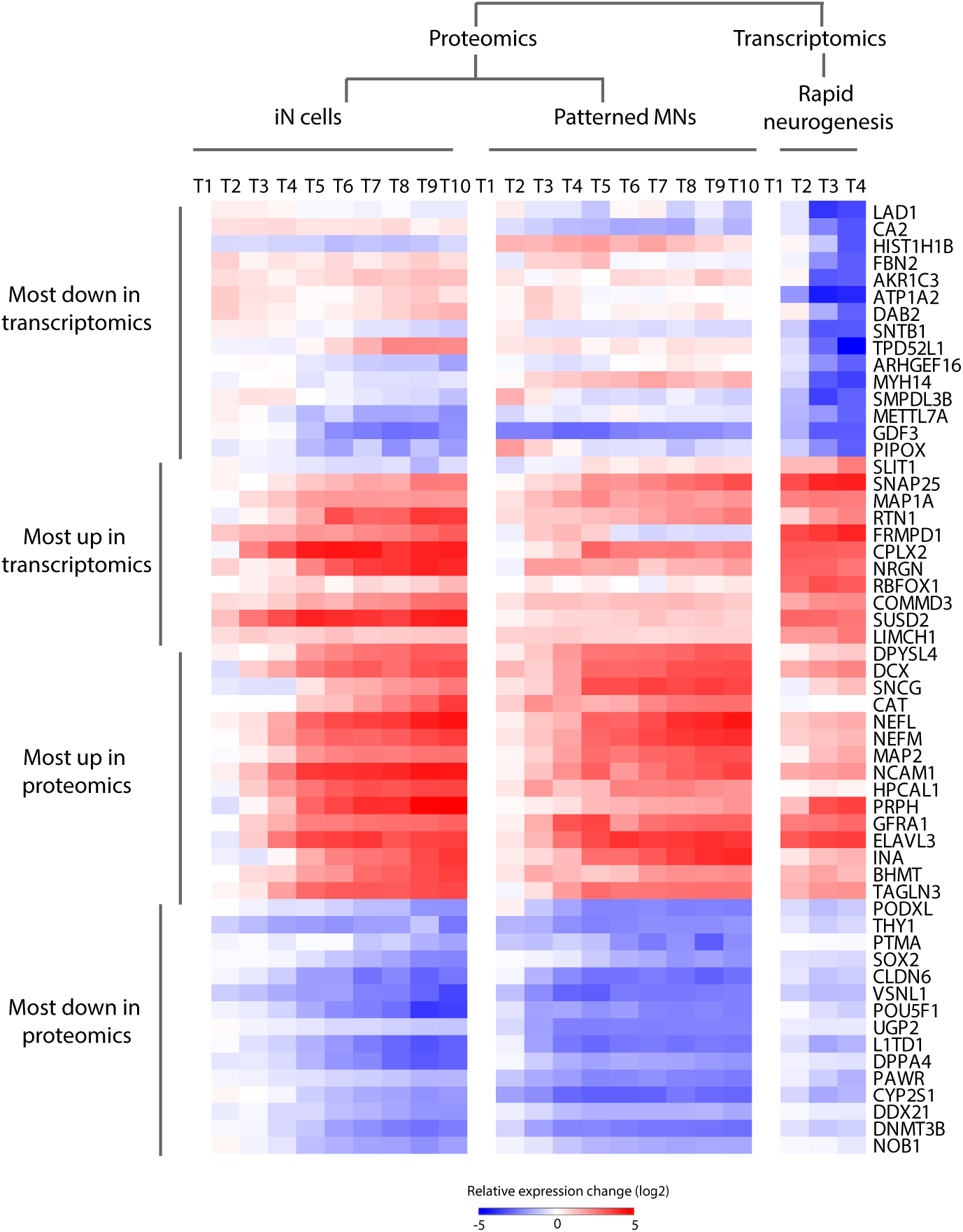
Comparison proteomic and transcriptomic profile. Heatmap of the top 15 most up- and downregulated proteins and top 15 most up- and downregulated genes along the course of differentiation towards neurons.

## DISCUSSION

Our study provide insight into the remodeling of the proteome during human neuronal development. To our knowledge, this is the first comprehensive proteomic profiling of human iPSC differentiation towards neurons. The rapid neurogenesis through transcriptional activation towards iN cells and the introduction of small molecules towards patterned MNs allowed us to identify key regulatory proteins involved in differentiation. Quantitative proteomics was applied to profile the dynamic changes of proteins during neuronal differentiation by using TMT 10-plex labeling coupled to high resolution LC-MS/MS, which resulted in the identification of 7230 proteins. We found a proteome-wide change in expression, which occurs in a two-step fashion, revealing a clear switch in protein expression levels, halfway our time window, leading to an anti-correlation between iPSC (T1) and neurons (T10). A similar but opposite trend was observed previously in a study monitoring proteome changes during the course of fibroblast reprogramming to iPSC [34]. According to GO classification, most of the proteins that downregulate during differentiation are located in the nucleus and are involved in RNA and DNA binding. Proteins that upregulate are mainly cytoskeletal proteins involved in cell communication and transport that could play a role in dendritic/axon outgrowth and branching. These findings are in line with our previous study, characterizing the proteome changes of rat hippocampal neuron development [35]. The majority of the proteins are steadily upregulated or downregulated along the course of differentiation. We highlighted the most extreme comparison of iPSC (T1) against mature neurons (T10) and showed the top 15 most up- or downregulated proteins during neuronal differentiation, consisting of proteins such as MAP2, a neuron-specific cytoskeletal protein having an established role in neurodevelopment [100]. Moreover, we detect several proteins (e.g. INA, DPYSL4, and RPS19) being highly upregulated with less established association with neuronal development suggesting that they may be considered as novel pan neuronal markers. In conjunction, we captured all TFs and cytoskeletal proteins in our data and illustrated their expression changes during neuronal differentiation towards the two subtypes. Several of these TF’s (PHC2, CRTC1, NCOA3 and SMAD3) have been reported to promote neuronal differentiation in zebrafish and mice, however their association to human neuronal differentiation was not detected. In addition, we illustrated several subunits of a protein family having differential expression profiles in both cell types, such as strong upregulation of TAGLN3 in both cell types and the upregulation of TAGLN and TAGLN2 only in iN cells. TAGLN was previously shown to regulate cytoskeleton organization, [101] and was differentially expressed in neuronal subpopulations of the rat central nervous system [102], highlighting this protein as an interesting target for further investigation. Furthermore, we provide a rich source of information of proteins associated with several signaling pathways, such as Wnt and Notch, involved in neuronal development. We identified several proteins associated to these signaling pathways being upregulated towards one of the two neuronal subtypes and being involved in cell fate pattering and development, further emphasizing their critical role in neuronal (subtype) differentiation. Also, we combined our proteomics data with comparable transcriptomic data during neuronal differentiation showing similarities as well as differences for several transcripts/proteins. These variations might be regulated by additional mechanisms such as posttranscriptional modifications that control the expression level of several proteins. In summary, we provide a quantitative view of key proteins that promote the loss of pluripotency and rapid neuronal development along the course of differentiation. This data constitute a rich resource that can be used in the future to better understand specific molecular mechanisms involved in neurodevelopment.

## Material and Methods

### Cell culture

#### iPSC generation

Generation of iPSC was performed using a previously established protocol [103]. Briefly, skin biopsies from healthy individuals were taken and fibroblast were maintained in mouse embryonic fibroblast (MEF) medium containing DMEM GlutaMAX (Life Technologies), 10% fetal bovine serum (Sigma Aldrich) and 1% penicillin/streptomycin (Life Technologies). The iPSC were generated by lentiviral transduction expressing OCT4, KLF4, SOX2 and c-MYC in MEF medium containing 4 mg/mL hexadimethrine bromide (Sigma). After 24 h of incubation, cells were cultured in MEF medium for another 5 days. Subsequently, cells were detached with Trypsin-EDTA (Life Technologies) and cultured in a 10 cm dish containing irradiated MEFs in human embryonic stem cell (huES) medium containing DMEM-F12 (Life Technologies), knockout 10% serum replacement (Life Technologies), 1% penicillin/streptomucin (Life Technologies), 2% L-glutamine (Life Technologies), 0.1% β-mercaptoethanol (Meck Millipore) and 20 ng/mL recombinant human fibroblast growth factor-basic (Life Technologies). After 3 to 6 weeks, colonies were picked manually and maintained in huES medium on irradiated MEFs for another 3-6 weeks. The iPSC were passaged using Accutase (Innovative Cell Technologies) and cultured feeder-free on Getrex (Life Technologies) coated dishes in mTeSR1 medium (STEMCELL technologies).

#### Glial cells

Newborn mouse forebrains homogenates were incubated with trypsin for 5 min at 37 °C. Following harsh trituration, cells were plated in MEM Alpha medium containing diglucose 20%, FBS 10%, Pen-strep 1%. After three passages, glial cells were used for co-culture purposes.

#### Virus generation

The two lentivirus were produced as described [104]. HEK293T cells plated on a 500 cm^2^ dish were cotransfected using MAXPEI solution with 35 μg pMD2.G, 65 μg psPAX and 100 μg pSIN-FUW-TeTO-Ngn2-P2A-EGFP-T2A-Puromycin or pSIN-FUW-M2rtTA in OPTI-MEM (Thermo Fisher Scientific). Six hours after transfection, the medium was replaced with OPTI-MEM supplemented with Penicillin-Streptomycin 1% (Thermo Fisher Scientific). The lentiviral particles were harvested 48 h after transfection, concentrated by tangential flow filtration using a 100 kDa cut-off spin filter (Millipore) and resuspended in phosphate buffered saline (PBS).

#### Differentiation

Generation of iN cells was performed using a slightly modified version of the protocol described in Yingsha Zhang *et al*, 2013 [19]. In short, on day 0, iPSC were treated with Accutase and plated at a density of 50 × 10^3^ cells/well of a 24-wells plate on top of a Matrigel (BD Biosciences) coated coverslip in mTeSR containing Y27632 10 μM (Miltenyibiotec). On day 1, medium was changed to mTeSR supplemented with 2,5 μl lentivirus and polybrene 8 μg/μL (Sigma-Aldrich). On day 2, medium was replaced with N2/DMEM/F12/NEAA (Invitrogen) supplemented with BDNF 10 μg/L (R&D systems), human NT-3 10 μg/L (ReproTech), mouse laminin 0.2 mg/L (Invitrogen) and doxycycline 2g/L (Clontech). On day 3, puromycin 1mg/L (Sigma-Aldrich) was added to the cells. On day 4, coverslips with neurons was transferred on top of a 12-well plate cultured glial cells in Neurobasal medium containing B27 (ThermoFisher), Glutamax (Invitrogen), BDNF, NT-3, doxycycline, Ara-C 2 g/L (Sigma-Aldrich). Every other day thereafter, 50% of the medium was changed. At predetermined time points, cells were fixed for immunohistochemistry or for proteomics approaches.

Patterned MN differentiation was performed using a slightly modified version of previously described protocol in Yves Maury *et al*, Nature biotechnology,2015 [20]. Briefly, on day 0, iPSC were dissociated with Accutase and resuspended in differentiation medium containing DMEM F-12, Neurobasal *(v/v)*, N2 supplement (Life Technologies), B27 without vitamin A (Life Technologies), Pen-strep 1%, ascorbic acid (0.5 μM, Sigma-Aldrich) and 5 μM Y27632 (STemGent). Embryoid body (EB) formation was accompanied by a standardized microwell assay [105]. IPSC were seeded at a density of 150 cells/microwell in differentiation medium. Chir-99021 (Tocris), LDN193189, SB-431542, Smoothened agonist (SAG, Calbiochem), retinoic acid (RA, Sigma-Aldrich), DAPT (Tocris), BDNF (Peprotech) and GDNF (Peprotech) were added at indicated time points and concentrations. Medium was changed every other day. After 2-3 days, EBs were flushed out of the microwell and transferred to a non-adherent 10 cm petri dish (Greiner Bio-one). On day 15, EBs were dissociated into single cells using Papain (Worthington Biochemical Corporation) and DNAse (Worthington Biochemical Corporation). Cells were plated on PDL (20 μg/mL, Sigma-Aldrich) and laminin (5 μg/mL, Invitrogen) coated coverslips at 60-70 % confluency.

#### Immunohistochemistry

At the time points of interest, cells were fixed in 4% paraformaldehyde in PBS for 20 min at room temperature and washed with PBS. Cells were blocked for 45 min at room temperature in blocking solution (0.1% Tween in PBS containing 2% bovine serum albumin (BSA) and 20 % goat serum). Cells were washed three times in PBS and primary antibody was incubated for 1 h at room temperature. After washing the cells three times, secondary antibody was applied and incubated for 1 h at room temperature in darkness. Primary and secondary antibodies were mixed in staining solution (0.1% Tween in PBS containing 5% goat serum). Cells were washed again and fixed with prolong gold and mounted on glass slides. The following commercial antibodies were used: rabbit anti-Tubbulin-β3 (Sigma), mouse anti-Isl-1 (DSHB), Hoechst or DAPI (invitrogen) rabbit-anti-FOXG1 (Abcam), mouse anti-TuJ1 (Covance). Alexa-488, and Alexa 546 conjugated secondary antibodies were obtained from Invitrogen.

#### Sample preparation for MS analysis

Samples were collected at indicated time points for MS analysis as follows. Cells were lysed in lysis buffer containing 8 M urea, 50 mM ammonium bicarbonate and one complete mini protease inhibitor (Roche). The lysate was sonicated on ice using Bioruptor-Diagenode and centrifuged at 2500 g for 10 min at 4 °C. Supernatant with the proteins was reduced with 4 mM dithiothreitol at 56 °C for 30 min and alkylated with 8 mM iodoacetamide at room temperature for 30 min in darkness. The lysate was enzymatically predigested with Lys-C (1:75; Wako) incubation for 4 h at 37 °C. The mixture was fourfold diluted with ammonium bicarbonate and digested with trypsin (1:100; Promega) at 37 °C. The sample was quenched by acidification with formic acid (FA, final concentration 10%) and peptides were desalted using a Sep-Pak C18 column (Waters). Peptides were dried in vacuum and resuspended in 50 mM triethyl ammonium bicarbonate at a final concentration of 5 mg/mL.

#### Tandem Mass Tag (TMT) 10-plex labelling

Aliquots of ~ 100 μg of each sample were chemically labeled with TMT reagents (Thermo Fisher). Peptides were resuspended in 80 μl resuspension buffer containing 50 mM HEPES buffer and 12.5 % acetonitrile (ACN, pH 8.5). TMT reagents (0.8 mg) were dissolved in 80 μl anhydrous ACN of which 20 μl was added to the peptides. Following incubation at room temperature for 1 hour, the reaction was quenched using 5% hydroxylamine in HEPES buffer for 15 min at room temperature. The TMT-labeled samples were pooled at equal protein ratios followed by vacuum centrifuge to near dryness and desalting using Sep-Pak C18 cartridges.

#### Off-line basic pH fractionation

We fractionated and pooled samples using basic pH Reverse Phase HPLC. Samples were solubilized in buffer A (5% ACN, 10 mM ammonium bicarbonate, pH 8.0) and subjected to a 50 min linear gradient from 18% to 45% ACN in 10 mM ammonium bicarbonate pH 8 at flow rate of 0.8 ml/min. We used an Agilent 1100 pump equipped with a degasser and a photodiode array (PDA) detector and Agilent 300 Extend C18 column (5 μm particles, 4.6 mm i.d., and 20 cm in length). The peptide mixture was fractionated into 96 fractions and consolidated into 24. Samples were acidified with 10% formic acid and vacuum-dried followed by redissolving with 5% formic acid/5% ACN for LC-MS/MS processing.

#### Mass spectrometry analysis

Mass spectrometry was performed on an Orbitrap Fusion (Thermo Fisher Scientific) and Orbitrap Fusion Lumos (Thermo Fisher Scientific) coupled to an Agilent 1290 HPLC system (Agilent Technologies). Peptides were separated on a double frit trap column of 20 mm x 100 μm inner diameter (ReproSil C18, Dr Maisch GmbH, Ammerbuch, Germany). This was followed by a 40 cm x 50 μm inner diameter analytical column (ReproSil Pur C18-AQ (Dr Maisch GmbH, Ammerbuch, Germany). Both columns were packed in-house. Trapping was done at 5 μl/min in 0.1 M acetic acid in H2O for 10 min and the analytical separation was done at 100 nl/min for 2 h by increasing the concentration of 0.1 M acetic acid in 80% acetonitrile *(v/v)*. The instrument was operated in a data-dependent mode to automatically switch between MS and MS^2^ for patterned MNs or MS and MS^3^ for iN cells. Full-scan MS spectra were acquired in the Orbitrap from m/z 350-1500 with a resolution of 60,000 FHMW, automatic gain control (AGC) target of 200,000 and maximum injection time of 50 ms. For the MS^2^ analysis, the ten most intense precursors at a threshold above 5,000 were selected with an isolation window of 1.2 Th after accumulation to a target value of 30,000 (maximum injection time was 115 ms). Fragmentation was carried out using higher-energy collisional dissociation (HCD) with collision energy of 38% and activation time of 0.1 ms. Fragment ion analysis was performed on Orbitrap with resolution of 60,000 FHMW and a low mass cut-off setting of 120 Th. For the MS^3^ analysis, first MS^2^ analysis was performed with CID fragmentation on top 10 most intense ions with AGC target of 10 000 and isolation window of 0.7 Th, followed by MS^3^ scan for each MS^2^ scan with HCD fragmentation with 35% collision energy. The MS isolation window was set to 2 m/z and AGC target to 100,000 and maximum injection time was 120 ms.

#### Data processing

Mass spectra were processed using Proteome Discover (version 2.1, Thermo Scientific). Peak list was searched using Swissprot database (version 2014_08) with the search engine Mascot (version 2.3, Matrix Science). The following parameters were used. The enzyme was specified as trypsin and allowed up to two missed cleavages. Taxonomy was chosen for Homo sapiens and precursor mass tolerance was set to 50 ppm with 0.05 Da fragment mass tolerance for MS^2^ analysis or 0.6 Da for MS^3^ analysis. TMT tags on lysine residues and peptide N termini and oxidation of methionine residues were set as dynamic modifications, while carbamidomethylation on cysteine residues was set as static modification. For the reporter ion quantification, integration tolerance was set to 20 ppm with the most confident centroid method. Results were filtered with a Mascot score of at least 20 and Percolator was used to adjust the peptide-spectrum matches (PSMs) to a false discovery rate (FDR) below 1%. Finally, peptides lower than 6 amino-acid residues were discarded.

#### Data visualization

The software Perseus was used to generate the plots, heatmaps and to calculate the Pearson correlation. For quantitative analysis, each time point was divided by T1, within each biological replicate. The ratios of the biological replicates were then log2 transformed. All the peptide ratios were normalized against the median. Volcano plots for each time point was generated and up- or downregulated proteins were considered significant with FDR < 0.05. Z scores were used to generate heatmaps. Functional analysis to enrich to GO, MF and CO terms were done using Gprox open source software package [106]. Clustering parameters such as Fuzzification, regulation threshold and number of clusters were set to 2.00, upper limit =-0.58 lower limit = 0.58 and 3, respectively. For the identification of time specific proteins between rostral and caudal neurons, volcano plots was generated comparing the two groups for each time point and proteins were considered to be significant with a fold change cut-off ≥ 1.5 and a p value ≤ 0.1. Furthermore, protein classification was performed using PANTHER [107] classification system and GO analysis and classification of transcription factors, cytoskeletal proteins were done using DAVID database. In addition, Reactome pathway database analysis was used to identify pathways enriched in several clusters.

#### Data availability

All mass spectrometry proteomics data have been deposited to the ProteomeXchange Consortium via the PRIDE partner repository with the dataset identifier PXD013303

## Acknowledgements

This work was supported by the Netherlands Organization for Scientific Research (NWO) through a VIDI grant for M.A. (723.012.102) and Proteins@Work, a program of the National Roadmap Large-scale Research Facilities of the Netherlands (project number 184.032.201). We thank the MIND facility at the UMC Utrecht for the access and help with the cell culture.

## Supplementary Information

**Figure S1.**
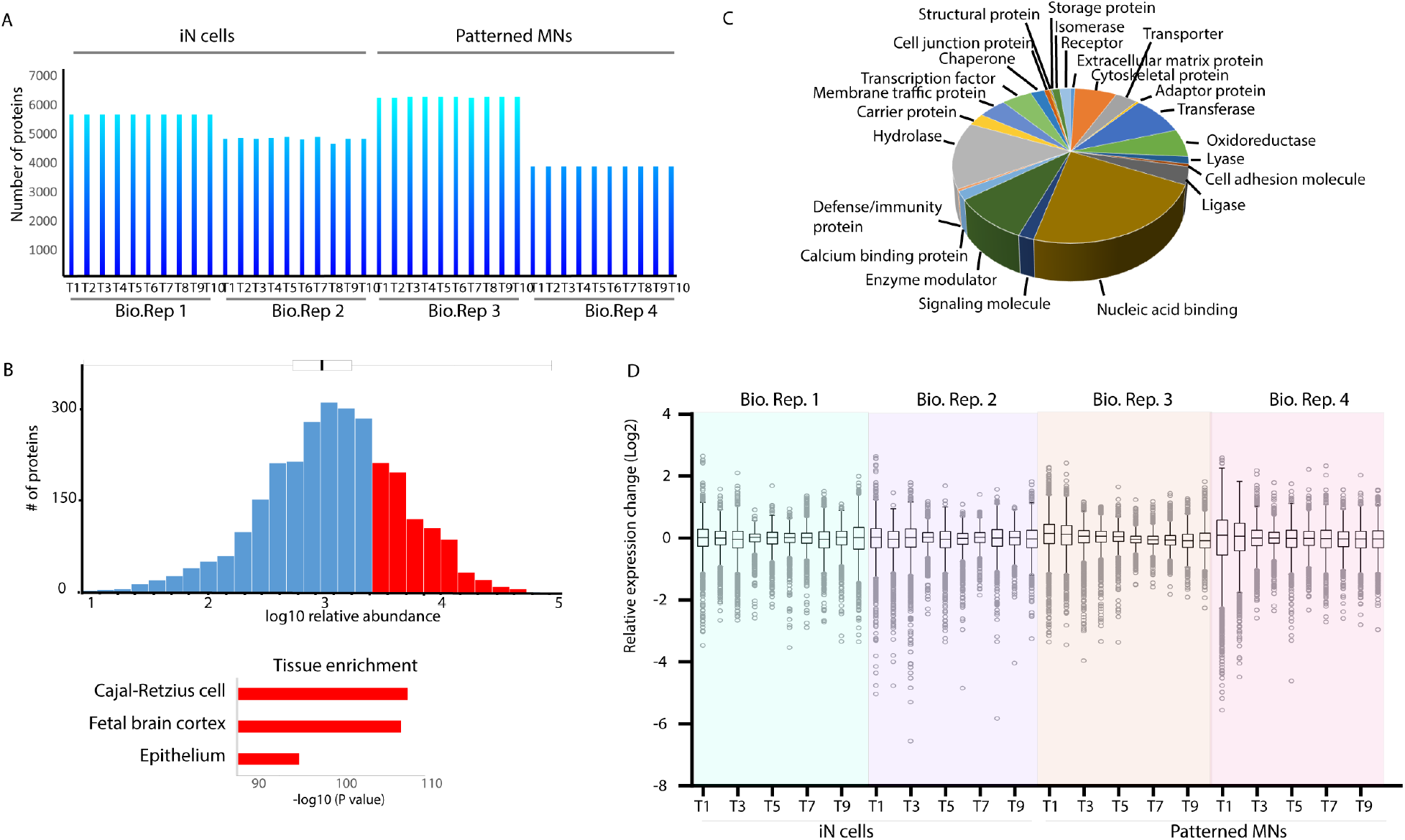
Global proteome analysis. (A) Bar graph showing the number of proteins identified in each biological replicate and in each time point. (B) Distribution of quantified protein abundances in all biological replicas and all time points, spanning four orders of magnitude. In red tissue enrichment analysis using DAVID, of the 25% most abundant proteins. (C). Pie charts display the distribution of protein classes, based on PANTHER classification from all quantified proteins. (D) Box plots of log2 transformed data normalized on protein peak areas of each individual biological replicate.

**Figure S2.**
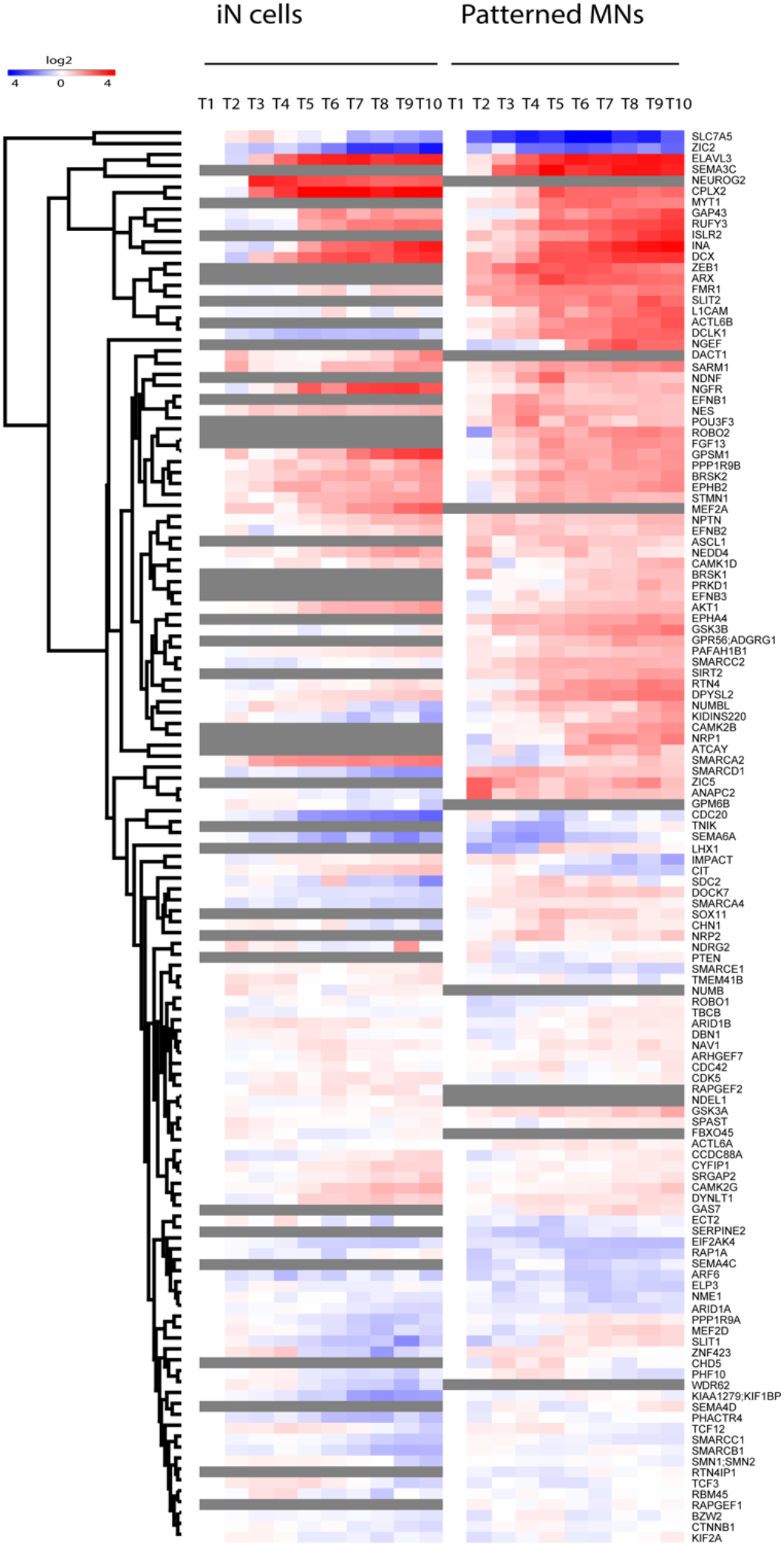
Neurogenesis. Neurogenesis associated proteins enriched using the Database for Annotation, Visualization, and Integrated Discovery (DAVID).

**Figure S3.**
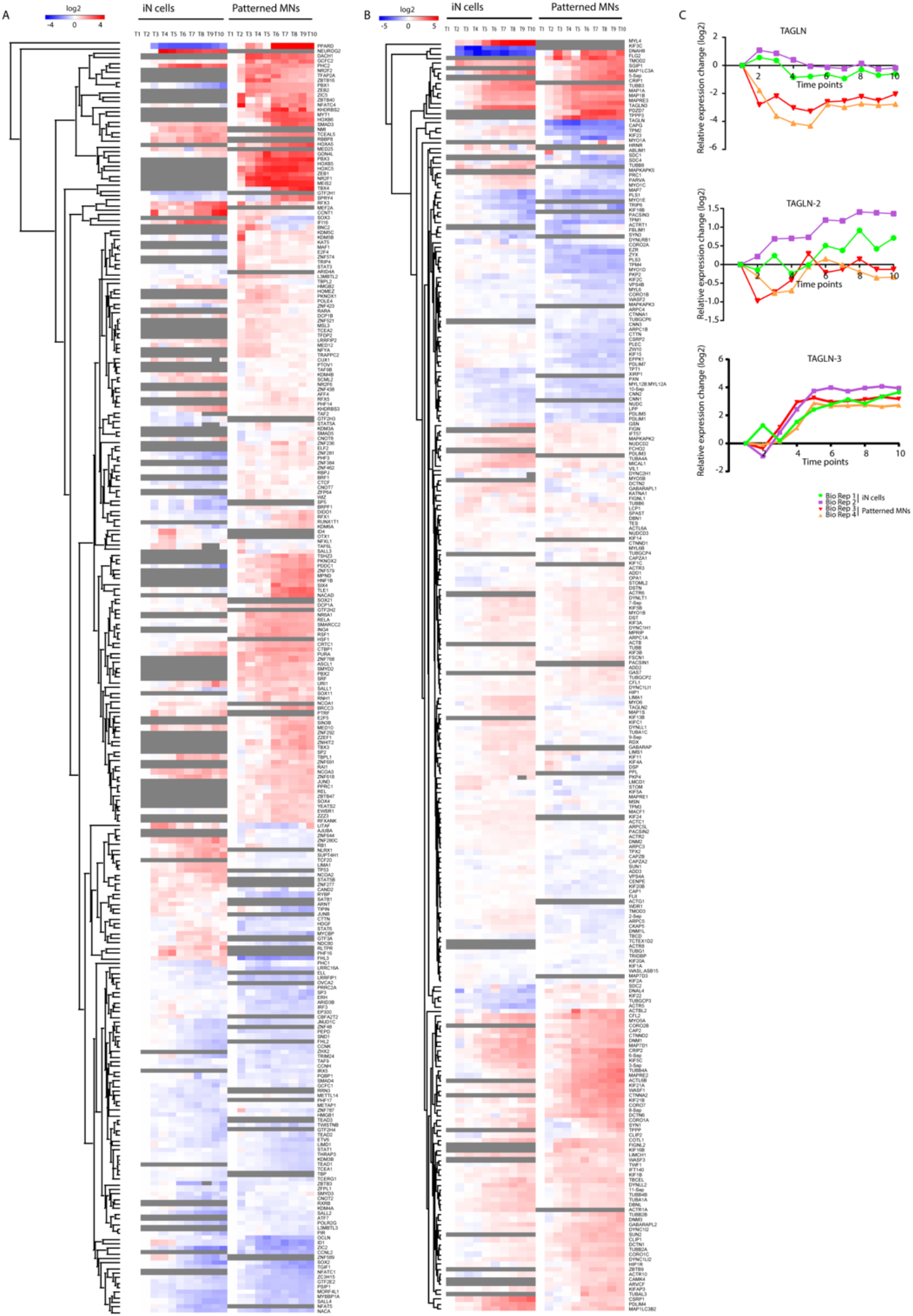
Transcription factors and cytoskeletal proteins. (A) Transcrioption factors enriched using the Database for Annotation, Visualization, and Integrated Discovery (DAVID). (B) Cytoskeletal proteins enriched using DAVID. (C) Example of the Transgelin family that undergo different expression during differentiation towards rostral and caudal neurons.

**Figure S4.**
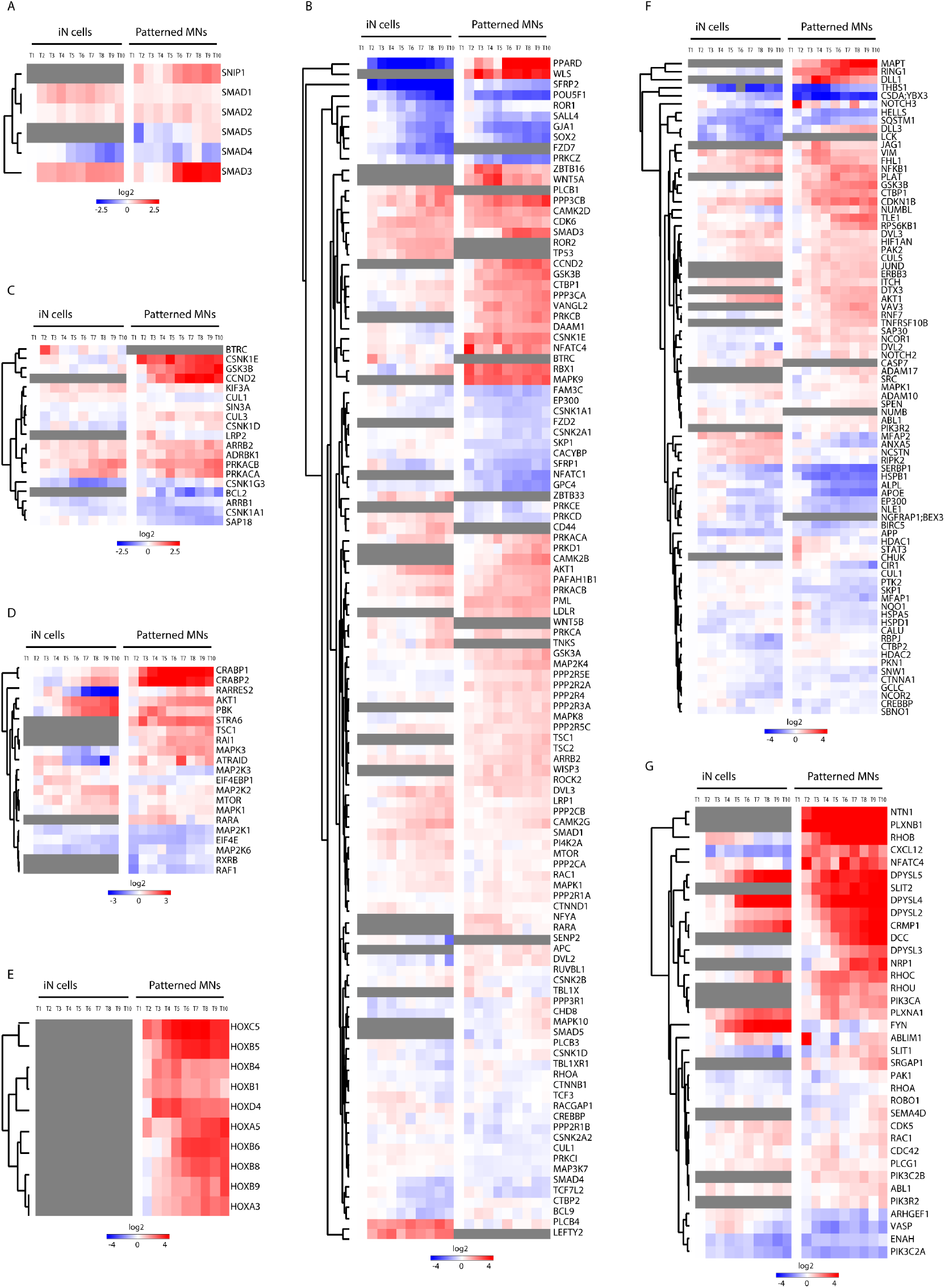
Signaling pathways. (A). Proteins related to TGFβ signaling enriched from KEGG and WIKI Pathways. (B). Proteins related to Wnt signaling. (C). Proteins related to Hedgehog signaling. (D). Proteins related to RA signaling. (E). Homeobox proteins all expressed in patterned MNs and none of the different members expressed in iN cells. (F). Proteins related to Notch signaling. (G). Proteins related to axon guidance

**Figure S5.**
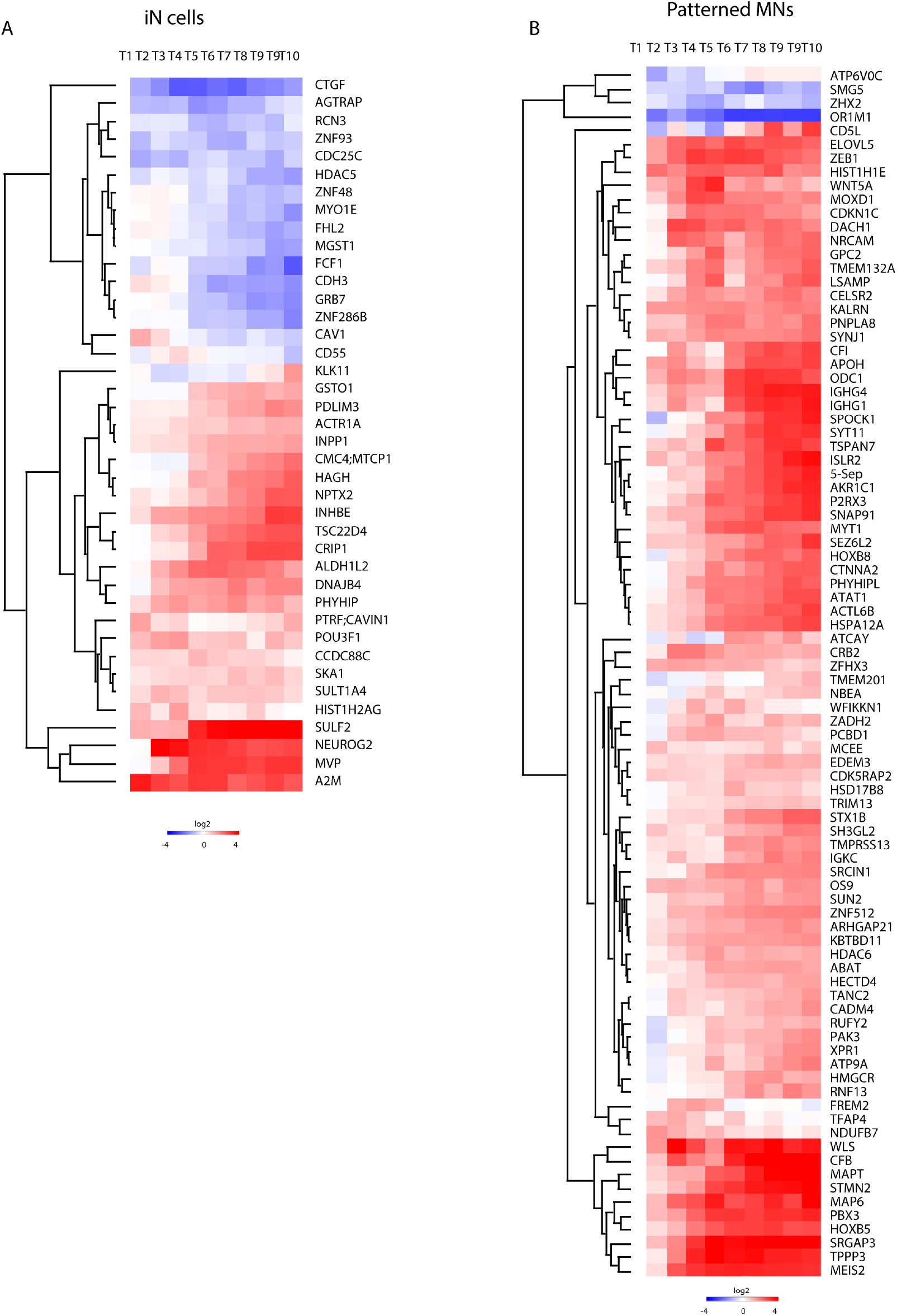
Proteins only identified in iN cells or patterned MNs. Proteins exclusively identified in (A) iN cells or (B) patterned MN differentiation. Proteins were selected for a twofold change in the two biological replicates in each group.

## REFERENCES

1. Herculano-Houzel, S., The human brain in numbers: a linearly scaled-up primate brain. Front Hum Neurosci, 2009. 3: p. 31.

2. Rockland, K.S., Non-uniformity of extrinsic connections and columnar organization. J Neurocytol, 2002. 31(3-5): p. 247–53.

3. Toro, R., et al., Key role for gene dosage and synaptic homeostasis in autism spectrum disorders. Trends Genet, 2010. 26(8): p. 363–72.

4. Hall, J., et al., Associative learning and the genetics of schizophrenia. Trends Neurosci, 2009. 32(6): p. 359–65.

5. Harvey, M., P. Belleau, and N. Barden, Gene interactions in depression: pathways out of darkness. Trends Genet, 2007. 23(11): p. 547–56.

6. Willsey, A.J., et al., Coexpression networks implicate human midfetal deep cortical projection neurons in the pathogenesis of autism. Cell, 2013. 155(5): p. 997–1007.

7. Parikshak, N.N., et al., Integrative functional genomic analyses implicate specific molecular pathways and circuits in autism. Cell, 2013. 155(5): p. 1008–21.

8. Takahashi, K., et al., Induction of pluripotent stem cells from adult human fibroblasts by defined factors. Cell, 2007. 131(5): p. 861–72.

9. Han, S.S., L.A. Williams, and K.C. Eggan, Constructing and deconstructing stem cell models of neurological disease. Neuron, 2011. 70(4): p. 626–44.

10. Marchetto, M.C. and F.H. Gage, Modeling brain disease in a dish: really? Cell Stem Cell, 2012. 10(6): p. 642–5.

11. Tao, Y. and S.C. Zhang, Neural Subtype Specification from Human Pluripotent Stem Cells. Cell Stem Cell, 2016. 19(5): p. 573–586.

12. Bertrand, N., D.S. Castro, and F. Guillemot, Proneural genes and the specification of neural cell types. Nat Rev Neurosci, 2002. 3(7): p. 517–30.

13. Kelava, I. and M.A. Lancaster, Stem Cell Models of Human Brain Development. Cell Stem Cell, 2016. 18(6): p. 736–48.

14. Busskamp, V., et al., Rapid neurogenesis through transcriptional activation in human stem cells. Mol Syst Biol, 2014. 10: p. 760.

15. Hoffmann, A., M. Ziller, and D. Spengler, Childhood-Onset Schizophrenia: Insights from Induced Pluripotent Stem Cells. Int J Mol Sci, 2018. 19(12).

16. Taoufik, E., et al., Synaptic dysfunction in neurodegenerative and neurodevelopmental diseases: an overview of induced pluripotent stem-cell-based disease models. Open Biol, 2018. 8(9).

17. Guo, W., et al., Current Advances and Limitations in Modeling ALS/FTD in a Dish Using Induced Pluripotent Stem Cells. Front Neurosci, 2017. 11: p. 671.

18. Sances, S., et al., Modeling ALS with motor neurons derived from human induced pluripotent stem cells. Nat Neurosci, 2016. 19(4): p. 542–53.

19. Zhang, Y., et al., Rapid single-step induction of functional neurons from human pluripotent stem cells. Neuron, 2013. 78(5): p. 785–98.

20. Maury, Y., et al., Combinatorial analysis of developmental cues efficiently converts human pluripotent stem cells into multiple neuronal subtypes. Nat Biotechnol, 2015. 33(1): p. 89–96.

21. Bielle, F., et al., Multiple origins of Cajal-Retzius cells at the borders of the developing pallium. Nat Neurosci, 2005. 8(8): p. 1002–12.

22. Rosskothen-Kuhl, N. and R.B. Illing, Gap43 transcription modulation in the adult brain depends on sensory activity and synaptic cooperation. PLoS One, 2014. 9(3): p. e92624.

23. Hoffman, P.N. and R.J. Lasek, The slow component of axonal transport. Identification of major structural polypeptides of the axon and their generality among mammalian neurons. J Cell Biol, 1975. 66(2): p. 351–66.

24. Lee, H.K., et al., Dynamic Ca2+-dependent stimulation of vesicle fusion by membrane-anchored synaptotagmin 1. Science, 2010. 328(5979): p. 760–3.

25. Graham, V., et al., SOX2 functions to maintain neural progenitor identity. Neuron, 2003. 39(5): p. 749–65.

26. Zeineddine, D., et al., The Oct4 protein: more than a magic stemness marker. Am J Stem Cells, 2014. 3(2): p. 74–82.

27. Mazzoni, E.O., et al., Synergistic binding of transcription factors to cell-specific enhancers programs motor neuron identity. Nat Neurosci, 2013. 16(9): p. 1219–27.

28. Li, X., et al., Small-Molecule-Driven Direct Reprogramming of Mouse Fibroblasts into Functional Neurons. Cell Stem Cell, 2015. 17(2): p. 195–203.

29. Vincent, A.J., P.W. Lau, and A.J. Roskams, SPARC is expressed by macroglia and microglia in the developing and mature nervous system. Dev Dyn, 2008. 237(5): p. 1449–62.

30. Zhou, S., et al., Survivin Improves Reprogramming Efficiency of Human Neural Progenitors by Single Molecule OCT4. Stem cells international, 2016. 2016: p. 4729535–4729535.

31. Gongidi, V., et al., SPARC-like 1 regulates the terminal phase of radial glia-guided migration in the cerebral cortex. Neuron, 2004. 41(1): p. 57–69.

32. Harkin, L.F., et al., Distinct expression patterns for type II topoisomerases IIA and IIB in the early foetal human telencephalon. J Anat, 2016. 228(3): p. 452–63.

33. Schuurmans, C., et al., Sequential phases of cortical specification involve Neurogenin-dependent and -independent pathways. Embo j, 2004. 23(14): p. 2892–902.

34. Hansson, J., et al., Highly coordinated proteome dynamics during reprogramming of somatic cells to pluripotency. Cell Rep, 2012. 2(6): p. 1579–92.

35. Frese, C.K., et al., Quantitative Map of Proteome Dynamics during Neuronal Differentiation. Cell Rep, 2017. 18(6): p. 1527–1542.

36. Liao, M.-L., et al., Distribution patterns of the zebrafish neuronal intermediate filaments inaa and inab. Journal of Neuroscience Research, 2019. 97(2): p. 202–214.

37. Quach, T.T., et al., Collapsin response mediator protein 3 increases the dendritic arborization of hippocampal neurons. Mol Psychiatry, 2015. 20(9): p. 1027.

38. da Rocha Boeira, T., et al., Polymorphism Located in the Upstream Region of the RPS19 Gene (rs2305809) Is Associated With Cervical Cancer: A Case-control Study. J Cancer Prev, 2018. 23(3): p. 147–152.

39. Moore, B.W., A soluble protein characteristic of the nervous system. Biochem Biophys Res Commun, 1965. 19(6): p. 739–44.

40. Filipek, A., et al., Calcyclin--Ca(2+)-binding protein homologous to glial S-100 beta is present in neurones. Neuroreport, 1993. 4(4): p. 383–6.

41. Yamashita, N., et al., Distribution of a specific calcium-binding protein of the S100 protein family, S100A6 (calcyclin), in subpopulations of neurons and glial cells of the adult rat nervous system. J Comp Neurol, 1999. 404(2): p. 235–57.

42. Chan, W.Y., et al., Differential expression of S100 proteins in the developing human hippocampus and temporal cortex. Microsc Res Tech, 2003. 60(6): p. 600–13.

43. Girard, F., et al., The EF-hand Ca(2+)-binding protein super-family: a genome-wide analysis of gene expression patterns in the adult mouse brain. Neuroscience, 2015. 294: p. 116–55.

44. Hupe, M., et al., Gene expression profiles of brain endothelial cells during embryonic development at bulk and single-cell levels. Sci Signal, 2017. 10(487).

45. Chantha, S.C., et al., The MIDASIN and NOTCHLESS genes are essential for female gametophyte development in Arabidopsis thaliana. Physiol Mol Biol Plants, 2010. 16(1): p. 3–18.

46. Kalus, I., et al., Sulf1 and Sulf2 Differentially Modulate Heparan Sulfate Proteoglycan Sulfation during Postnatal Cerebellum Development: Evidence for Neuroprotective and Neurite Outgrowth Promoting Functions. PLoS One, 2015. 10(10): p. e0139853.

47. Kalus, I., et al., Differential involvement of the extracellular 6-O-endosulfatases Sulf1 and Sulf2 in brain development and neuronal and behavioural plasticity. J Cell Mol Med, 2009. 13(11–12): p. 4505–21.

48. Briscoe, J. and J. Ericson, Specification of neuronal fates in the ventral neural tube. Current Opinion in Neurobiology, 2001. 11(1): p. 43–49.

49. Couillard-Despres, S., et al., Doublecortin expression levels in adult brain reflect neurogenesis. Eur J Neurosci, 2005. 21(1): p. 1–14.

50. Kim, D.H., et al., Pan-neuronal calcium imaging with cellular resolution in freely swimming zebrafish. Nature Methods, 2017. 14: p. 1107.

51. Abbott, N.J., et al., Structure and function of the blood-brain barrier. Neurobiology of Disease, 2010. 37(1): p. 13–25.

52. Zhang, Y. and L. Niswander, Zic2 is required for enteric nervous system development and neurite outgrowth: a mouse model of enteric hyperplasia and dysplasia. Neurogastroenterology and motility: the official journal of the European Gastrointestinal Motility Society, 2013. 25(6): p. 538–541.

53. Morrison, S.J., Neuronal differentiation: proneural genes inhibit gliogenesis. Curr Biol, 2001. 11(9): p. R349–51.

54. Serre, A., et al., Overexpression of basic helix-loop-helix transcription factors enhances neuronal differentiation of fetal human neural progenitor cells in various ways. Stem Cells Dev, 2012. 21(4): p. 539–53.

55. Ladewig, J., et al., Small molecules enable highly efficient neuronal conversion of human fibroblasts. Nat Methods, 2012. 9(6): p. 575–8.

56. Conkright, M.D., et al., TORCs: transducers of regulated CREB activity. Mol Cell, 2003. 12(2): p. 413–23.

57. Li, S., et al., TORC1 regulates activity-dependent CREB-target gene transcription and dendritic growth of developing cortical neurons. J Neurosci, 2009. 29(8): p. 2334–43.

58. Mendioroz, M., et al., CRTC1 gene is differentially methylated in the human hippocampus in Alzheimer’s disease. Alzheimers Res Ther, 2016. 8(1): p. 15.

59. Storchel, P.H., et al., A large-scale functional screen identifies Nova1 and Ncoa3 as regulators of neuronal miRNA function. Embo j, 2015. 34(17): p. 2237–54.

60. Casari, A., et al., A Smad3 transgenic reporter reveals TGF-beta control of zebrafish spinal cord development. Developmental Biology, 2014. 396(1): p. 81–93.

61. Inoue, T., et al., Mouse Zic5 deficiency results in neural tube defects and hypoplasia of cephalic neural crest derivatives. Dev Biol, 2004. 270(1): p. 146–62.

62. Quadrato, G., et al., Nuclear factor of activated T cells (NFATc4) is required for BDNF-dependent survival of adult-born neurons and spatial memory formation in the hippocampus. Proceedings of the National Academy of Sciences, 2012. 109(23): p. E1499–E1508.

63. Mann, S.S. and J.A. Hammarback, Gene localization and developmental expression of light chain 3: a common subunit of microtubule-associated protein 1A(MAP1A) and MAP1B. J Neurosci Res, 1996. 43(5): p. 535–44.

64. Sullivan, K.F. and D.W. Cleveland, Identification of conserved isotype-defining variable region sequences for four vertebrate beta tubulin polypeptide classes. Proc Natl Acad Sci U S A, 1986. 83(12): p. 4327–31.

65. Mori, K., et al., Neuronal protein NP25 interacts with F-actin. Neurosci Res, 2004. 48(4): p. 439–46.

66. Stiles, J. and T.L. Jernigan, The basics of brain development. Neuropsychology review, 2010. 20(4): p. 327–348.

67. Tao, Y. and S.-C. Zhang, Neural Subtype Specification from Human Pluripotent Stem Cells. Cell stem cell, 2016. 19(5): p. 573–586.

68. Chambers, S.M., et al., Highly efficient neural conversion of human ES and iPS cells by dual inhibition of SMAD signaling. Nat Biotechnol, 2009. 27(3): p. 275–80.

69. Martin, B.L. and D. Kimelman, Wnt signaling and the evolution of embryonic posterior development. Curr Biol, 2009. 19(5): p. R215–9.

70. Ciani, L. and P.C. Salinas, WNTs in the vertebrate nervous system: from patterning to neuronal connectivity. Nat Rev Neurosci, 2005. 6(5): p. 351–62.

71. Budnik, V. and P.C. Salinas, Wnt signaling during synaptic development and plasticity. Curr Opin Neurobiol, 2011. 21(1): p. 151–9.

72. Hirabayashi, Y., et al., The Wnt/beta-catenin pathway directs neuronal differentiation of cortical neural precursor cells. Development, 2004. 131(12): p. 2791–801.

73. Nordstrom, U., et al., An early role for WNT signaling in specifying neural patterns of Cdx and Hox gene expression and motor neuron subtype identity. PLoS Biol, 2006. 4(8): p. e252.

74. Bhatt, P.M. and R. Malgor, Wnt5a: a player in the pathogenesis of atherosclerosis and other inflammatory disorders. Atherosclerosis, 2014. 237(1): p. 155–62.

75. Zhu, X., et al., Wls-mediated Wnts differentially regulate distal limb patterning and tissue morphogenesis. Dev Biol, 2012. 365(2): p. 328–38.

76. Barna, M., P.P. Pandolfi, and L. Niswander, Gli3 and Plzf cooperate in proximal limb patterning at early stages of limb development. Nature, 2005. 436(7048): p. 277–81.

77. Serfling, E., et al., NFATc1 autoregulation: A crucial step for cell-fate determination. Vol. 27. 2006. 461–9.

78. Yang, Y.R., et al., Primary phospholipase C and brain disorders. Adv Biol Regul, 2016. 61: p. 80–5.

79. Lara-Ramirez, R., E. Zieger, and M. Schubert, Retinoic acid signaling in spinal cord development. Int J Biochem Cell Biol, 2013. 45(7): p. 1302–13.

80. Cayuso, J., et al., The Sonic hedgehog pathway independently controls the patterning, proliferation and survival of neuroepithelial cells by regulating Gli activity. Development, 2006. 133(3): p. 517–28.

81. Numata-Uematsu, Y., et al., Inhibition of collapsin response mediator protein-2 phosphorylation ameliorates motor phenotype of ALS model mice expressing SOD1G93A. Neuroscience Research, 2018.

82. Kowalczyk, A., et al., The critical role of cyclin D2 in adult neurogenesis. J Cell Biol, 2004. 167(2): p. 209–13.

83. Chen, R., et al., Single-Cell RNA-Seq Reveals Hypothalamic Cell Diversity. Cell Reports, 2017. 18(13): p. 3227–3241.

84. Papalopulu, N. and C. Kintner, A posteriorising factor, retinoic acid, reveals that anteroposterior patterning controls the timing of neuronal differentiation in Xenopus neuroectoderm. Development, 1996. 122(11): p. 3409–18.

85. Elkabetz, Y., et al., Human ES cell-derived neural rosettes reveal a functionally distinct early neural stem cell stage. Genes Dev, 2008. 22(2): p. 152–65.

86. Borghese, L., et al., Inhibition of notch signaling in human embryonic stem cell-derived neural stem cells delays G1/S phase transition and accelerates neuronal differentiation in vitro and in vivo. Stem Cells, 2010. 28(5): p. 955–64.

87. Adams, J.C. and F.M. Watt, Regulation of development and differentiation by the extracellular matrix. Development, 1993. 117(4): p. 1183–98.

88. Christopherson, K.S., et al., Thrombospondins Are Astrocyte-Secreted Proteins that Promote CNS Synaptogenesis. Cell, 2005. 120(3): p. 421–433.

89. Rasmussen, J.P. and A. Sagasti, Learning to swim, again: Axon regeneration in fish. Exp Neurol, 2017. 287(Pt 3): p. 318–330.

90. Dickson, B.J., Molecular Mechanisms of Axon Guidance. Science, 2002. 298(5600): p. 1959–1964.

91. Bashaw, G.J., et al., Repulsive axon guidance: Abelson and Enabled play opposing roles downstream of the roundabout receptor. Cell, 2000. 101(7): p. 703–15.

92. Desmond, J.C., et al., The Aldo-Keto Reductase AKR1C3 Is a Novel Suppressor of Cell Differentiation That Provides a Plausible Target for the Non-Cyclooxygenase-dependent Antineoplastic Actions of Nonsteroidal Anti-Inflammatory Drugs. Cancer Research, 2003. 63(2): p. 505–512.

93. Capo-Chichi, C.D., et al., Alteration of Differentiation Potentials by Modulating GATA Transcription Factors in Murine Embryonic Stem Cells. Stem cells international, 2010. 2010: p. 602068–602068.

94. Hoerder-Suabedissen, A., et al., Expression profiling of mouse subplate reveals a dynamic gene network and disease association with autism and schizophrenia. Proceedings of the National Academy of Sciences of the United States of America, 2013. 110(9): p. 3555–3560.

95. Fu, X., et al., Doublecortin (Dcx) family proteins regulate filamentous actin structure in developing neurons. The Journal of neuroscience: the official journal of the Society for Neuroscience, 2013. 33(2): p. 709–721.

96. Ince-Dunn, G., et al., Neuronal Elav-like (Hu) proteins regulate RNA splicing and abundance to control glutamate levels and neuronal excitability. Neuron, 2012. 75(6): p. 1067–1080.

97. Fleming, M.S., et al., Cis and trans RET signaling control the survival and central projection growth of rapidly adapting mechanoreceptors. eLife, 2015. 4: p. e06828–e06828.

98. Jones, I., et al., Development and validation of an in vitro model system to study peripheral sensory neuron development and injury. Scientific reports, 2018. 8(1): p. 15961–15961.

99. Lin, M., et al., RNA-Seq of human neurons derived from iPS cells reveals candidate long noncoding RNAs involved in neurogenesis and neuropsychiatric disorders. PLoS One, 2011. 6(9): p. e23356.

100. Neve, R.L., et al., Identification of cDNA clones for the human microtubule-associated protein tau and chromosomal localization of the genes for tau and microtubule-associated protein 2. Brain Res, 1986. 387(3): p. 271–80.

101. Elsafadi, M., et al., Transgelin is a TGFβ-inducible gene that regulates osteoblastic and adipogenic differentiation of human skeletal stem cells through actin cytoskeleston organization. Cell death & disease, 2016. 7(8): p. e2321–e2321.

102. Ren, W.-Z., et al., The identification of NP25: a novel protein that is differentially expressed by neuronal subpopulations. Molecular Brain Research, 1994. 22(1): p. 173–185.

103. Harschnitz, O., et al., Autoantibody pathogenicity in a multifocal motor neuropathy induced pluripotent stem cell-derived model. Ann Neurol, 2016. 80(1): p. 71–88.

104. Naldini, L., et al., In vivo gene delivery and stable transduction of nondividing cells by a lentiviral vector. Science, 1996. 272(5259): p. 263–7.

105. Rivron, N.C., et al., Tissue deformation spatially modulates VEGF signaling and angiogenesis. Proc Natl Acad Sci U S A, 2012. 109(18): p. 6886–91.

106. Rigbolt, K.T., J.T. Vanselow, and B. Blagoev, GProX, a user-friendly platform for bioinformatics analysis and visualization of quantitative proteomics data. Mol Cell Proteomics, 2011. 10(8): p. O110.007450.

107. Mi, H., et al., PANTHER version 6: protein sequence and function evolution data with expanded representation of biological pathways. Nucleic Acids Res, 2007. 35(Database issue): p. D247–52.

